# Evidenced by Indigenous and Western Science: An Arctic Nation Building Project Threatens Caribou and Inuit Harvesting Rights

**DOI:** 10.64898/2026.04.16.718946

**Authors:** Andrea Hanke, Amanda Dumond, Susan Kutz, David Borish

## Abstract

Canada’s ambition for mineral security and its responsibilities to protect at-risk species and uphold Indigenous rights clash in the case of the Grays Bay Road and Port (GBRP) in Nunavut, an infrastructure project intended to unlock critical mineral deposits. We compiled Indigenous and Western science through a density analysis of caribou harvesting data near the proposed project site. We identified three consistently used harvesting hotspots, with the most significant hotspot lying directly in the path of the proposed GBRP project. These results indicate that the GBRP project will have significant and unmitigable negative effects on caribou conservation, food security, and Inuit harvesting rights. Prime Minister Carney claims that middle power countries must act consistently in this era of geopolitical rupture; this commitment must transfer to natural resource development reviews so that decision-making may be consistent and rooted in cross-legislation responsibilities and values, including the land claims agreements between Indigenous groups and the Government of Canada.

## Introduction

Geoeconomic confrontation (sanctions, tariffs, investment screening etc.) is the top short- and intermediate-term risk to global stability in 2026, a condition both causing, and a consequence of, weakened multilateral institutions (World Economic Forum, 2026). Canada is not a geoeconomic stronghold and must approach this new global order with strategy and purpose, stated as much during Canadian Prime Minister Mark Carney’s special address at the Davos World Economic Forum that called for middle-power countries to rally together and create coalitions that work today alongside building strong domestic economies (Carney, 2026). Canada, with the end game of bolstering their domestic economy and reducing vulnerability to international volatility, has created a new critical mineral strategy that prioritizes growing Canadian expertise across the mineral supply chain (Government of Canada, 2022; Kalantzakos, 2020).

Alongside establishing new critical mineral coalitions, Canada began listing development projects of national interest for streamlined federal regulatory approval, including those for mineral security (One Canadian Economy Act, 2025). One project considered for review is the Grays Bay Port and Road (GBRP) project, the northern section of the Arctic Economic and Security Corridor (AESC) which is planned to connect Yellowknife, Northwest Territories to a deep water port at Grays Bay, Nunavut on the Northwest Passage via an all-season road (Fig. 1; Peres & Stanley, 2025). The GBRP project is fundamentally an infrastructure project intended to unlock mineral mining in western Nunavut, including proposals for copper and zinc mines, and is currently under territorial review by the Nunavut Impact Review Board (Peres & Stanley, 2025).

**Figure 1.**
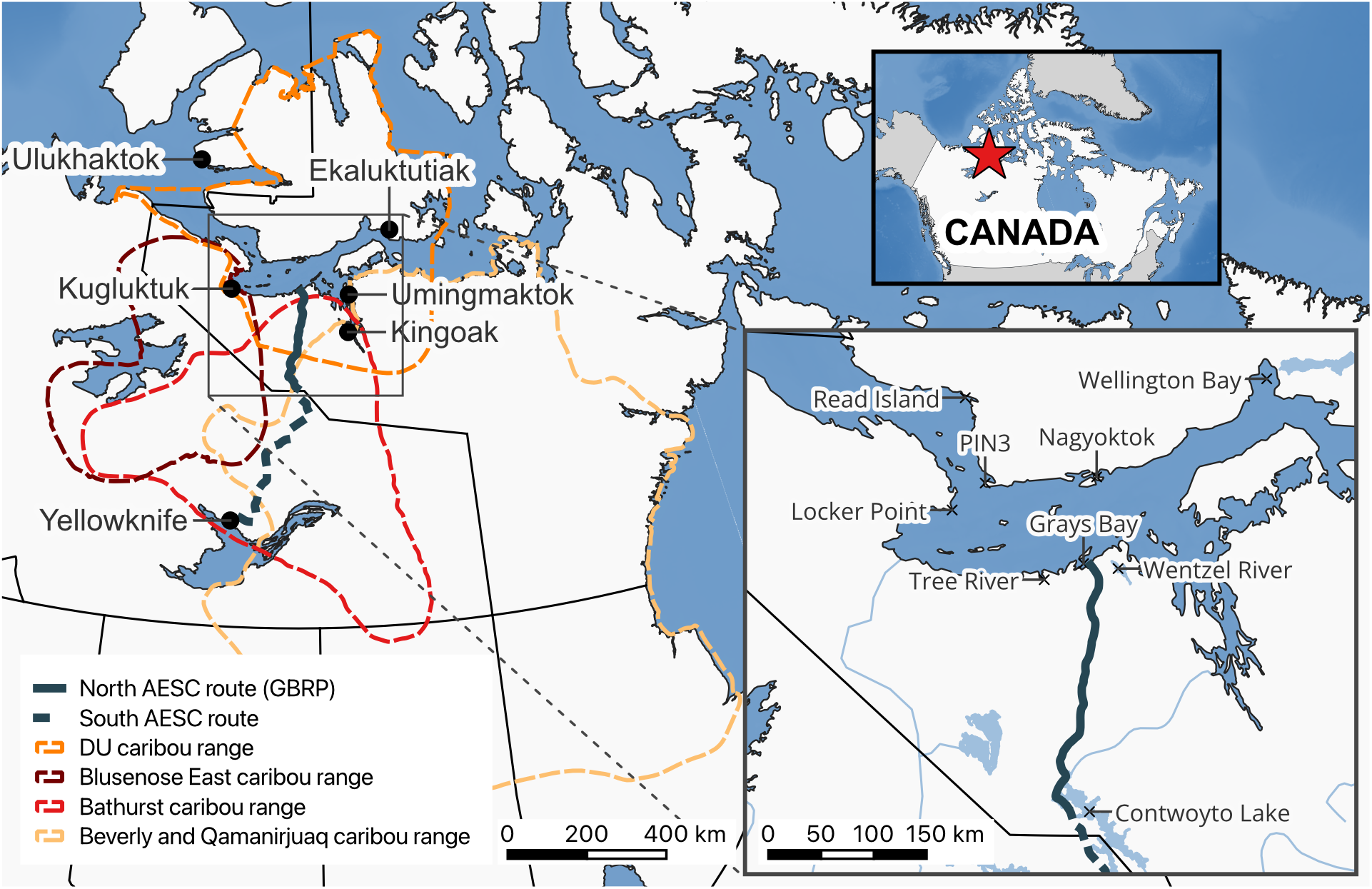
Project area and relevant caribou ranges. Figure includes the proposed route for the AESC, the caribou ranges for the DU, Bathurst, Beverly, and Qamanirjuaq herds (Beverly and Qamanirjuaq Caribou Management Board, 2023; Government of Northwest Territories, n.d., 2019; Worthington et al., 2018), locations of nearby communities, and a selection of placenames relevant for the project results.

Whether under streamlined federal review or territorial review, the Crown has a duty to consult and accommodate rightsbearing groups when a project could have adverse impacts on treaty rights and with the aim of securing free, prior, and informed consent (The Constitution Act, 1982). The GBRP project is under jurisdiction of three main rightsbearing groups under the Nunavut Act: Nunavut Tunngavik Incorporated and the Kitikmeot Inuit Association represent title and collective rights, while Hunters and Trappers Organizations represent Inuit harvesting rights (Nunavut Land Claims Agreement Act, 1993). Submissions to the Nunavut Impact Review Board document opposing views on the GBRP project, where the Kitikmeot Inuit Association (as a rights holder, previous project proponent, and major shareholder of the current proponent) wants the development to go forward while the Hunters and Trappers Organizations of Burnside and Kugluktuk strongly oppose the project due to concerns of irreversible negative impact to Inuit harvesting rights, particularly that of caribou, and local food security (Peres & Stanley, 2025).

Inuit harvesting rights are analogous with the Canadian right to religious freedom, and they are deeply steeped in how Inuit get food on the table for their families. Recent estimates place the Nunavut country food system at a worth of $198 million annually, clearly an incredibly important system in a territory that experiences a rate of food insecurity at 62.6% compared to 22.9% in the provinces (Statistics Canada, 2024; Warltier et al., 2021). Caribou are amongst the top consumed country foods in Nunavut, and they often provide the basis for Inuit culture and identity (Warltier et al., 2021).

The GBRP project location is home to four migratory caribou herds, three which are at-risk: Bathurst caribou (*Rangifer tarandus groenlandicus*) from spring through fall; Beverly caribou (*R. t. groenlandicus*) and Dolphin and Union (DU) caribou (*R. t. groenlandicus x pearyi*) during the winter and spring (Beverly and Qamanirjuaq Caribou Management Board, 2023; Government of Northwest Territories, 2019; Species at Risk Committee, 2023). In particular, the DU caribou are of utmost importance to the communities of Ulukhaktok and Paulatuk in Northwest Territories and Kugluktuk, Cambridge Bay, Bathurst Inlet, and Umingmaktok (Burnside) in Nunavut. The Committee on the Status of Endangered Wildlife in Canada reassessed the DU caribou herd as Endangered in 2017 (COSEWIC, 2017). Three years later, the Government of Nunavut enacted a total allowable harvest interim order to limit Nunavummiut harvesting to 42 DU caribou (Wildlife Act, 2020). The Nunavut Wildlife Management Board supported the federal uplisting in 2022 (Nunavut Wildlife Management Board, 2022), and the Government of Northwest Territories uplisted the herd to Endangered under their territorial Species at Risk Act in 2024 (Northwest Territories, 2024). Despite mounting support, the federal decision to uplist the DU caribou herd is still pending today.

## Methods

Our objective was to compile Indigenous and Western Knowledge to determine important locations for Inuit harvesting of DU caribou and to understand how that relates to the proposed GBRP project. We used a hexagonal binning, density analysis on harvesting data from three sources (see Fig. 1 for project location):

1. The Community-Based Muskox and Caribou Surveillance Program with the Kutz Research Group and Kugluktuk Angoniatit Association (140 latitude and longitude point data from 2018-2024). The primary data collection method is through harvester completed sample kits, where harvesters voluntarily collect biological samples from their catches, complete a form, and return the package to their local government wildlife officer or Hunters and Trappers organization to receive an honorarium. Research licensed under the Government of Nunavut, Wildlife Research Permit (WL 2024-041).
2. Department of Environment, Government of Nunavut (267 latitude and longitude point data from 2018-2024; mandatory harvest reporting since 2020). Data provided under a data-sharing agreement between the Government of Nunavut and Cloudberry Connections.
3. Indigenous Knowledge participatory mapping during 25 interviews with 33 residents of Kugluktuk in 2018-2020 (polygon data delineated by decade from 1980-2020). Expert caribou harvesters were invited to participate in the study based on recommendations by the Kugluktuk Angoniatit Association (purposive sampling) and suggestions of other harvesters from people already involved in the interviews (snowball sampling) (Hanke et al., 2024). Data collection was approved by Conjoint Faculties Research Ethics Board at the University of Calgary (REB17-2427) and licensed by the Nunavut Research Institute (04 013 23R-M).

To get the point and polygon data into similar formats, we broke the polygons up along a 10 km hexagon grid and created centroids for each new hexagon datum. Using the same 10 km hexagon grids, we counted the number of harvest reports and Indigenous Knowledge data points (latitude and longitude data points and proxy centroids) within each hexagon of the hexagon grid to get a total count of how many data points were within each 10 km hexagon area. Then, we symbolized each hexagon with a count of 1 or higher (removing the hexagons containing no data points).

## Results

The spatial data from harvest locations and participatory mapping covered 299,623 km^2^ of land (Table 1; Fig. 2) and clustered around three main harvest areas:

**Table 1.** Summary of spatial data and result hotspots.

| Feature | Area (km<sup>2</sup>) | # of data points | # of latitude and longitude data | # of interviews |
| --- | --- | --- | --- | --- |
| All data | 298,814 | 1-48 | 407 | 25 |
| All hotspots | 10,631 | 14-48 | 329 | 25 |
| Hotspot 1:<br>Tree River to Wentzel River | 4,738 | 14-48 | 233 | 25 |
| Hotspot 2:<br>Read Island, PIN3, Nagyoktok | 5,777 | 14-29 | 93 | 24 |
| Hotspot 3:<br>Locker Point | 116 | 15 | 3 | 12 |

**Figure 2.**
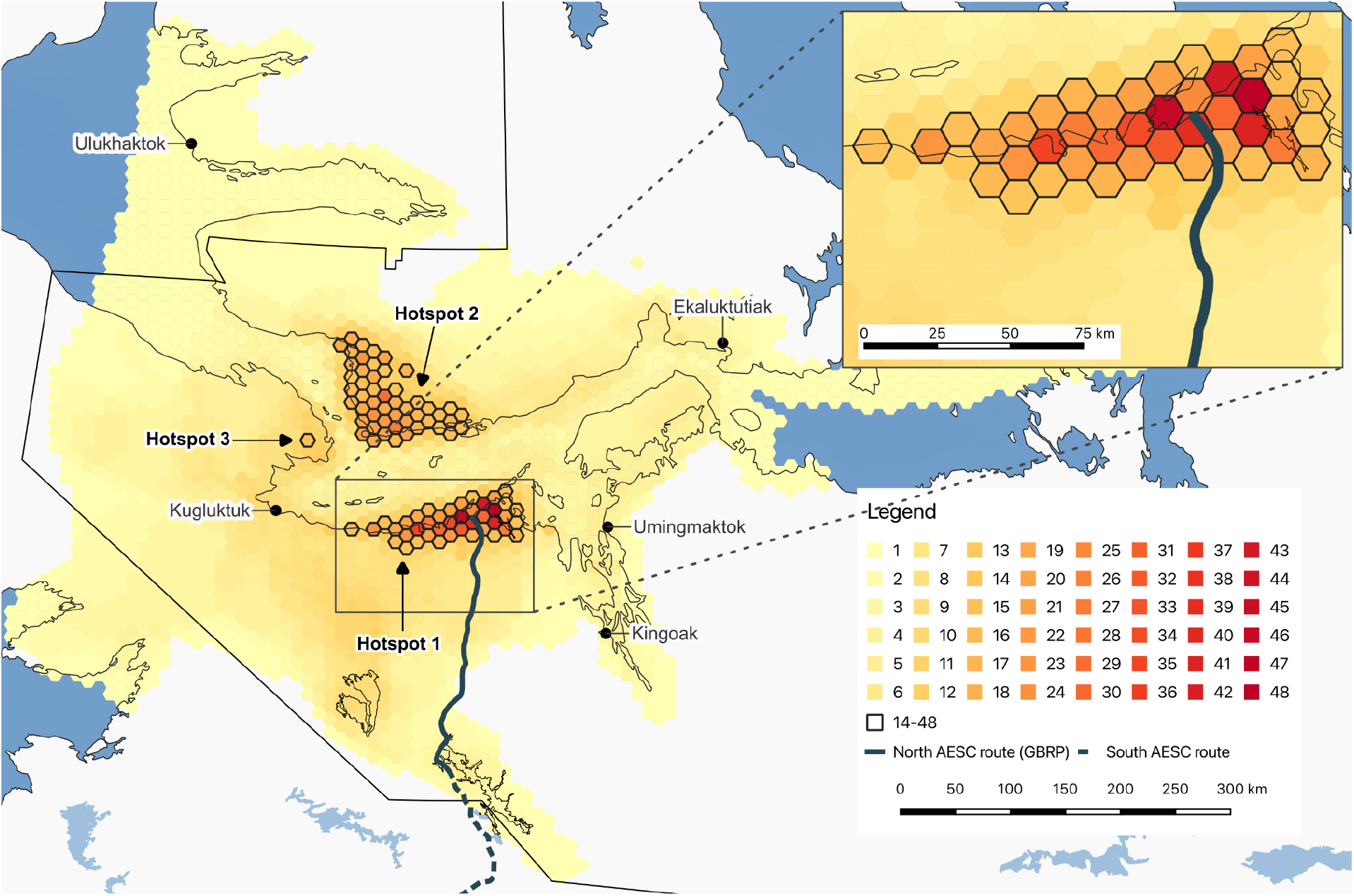
Density map of results that highlights three harvesting hotspots. Figure includes the proposed route for the AESC. Outlined hexagons have a data count (specific harvest locations and Traditional Knowledge estimates) between 14 and 48; all remaining hexagons have data counts between 1-13.

1. Between Tree River and Wentzel River, mostly when there is sea ice and mostly during April and May
2. Inshore from Read Island to PIN3 to Nagyoktok, mostly when there was no sea ice and mostly in August and September
3. Inshore from Locker Point

We reviewed changes in annual harvesting locations; our analysis showed no meaningful changes to annual harvesting locations between 2018-2024 within our data set.

The three identified hotspots are used consistently by caribou and for Inuit harvesting, and each one represents a key area for caribou survival and connection to Inuit cultural practices for a particular season. The largest and most dense hotspot between Tree River and Wentzel River lies directly in the path of the proposed GBRP project.

## Discussion

### Caribou cannot sustain the cumulative impacts of the GBRP project

Habitat protection is a key aspect of caribou conservation. Habitat alteration risks higher stress levels, restricted genetic flow, reduced resilience to predation and disease, higher susceptibility to rain-on-snow events, and a higher conservation risk (Dickie et al., 2023; Ewacha et al., 2017; Lesmerises et al., 2019; Lucas et al., 2019). Linear features alter caribou behaviour, facilitate increased predator hunting efficiency and presence, and tend to regenerate quite slowly, thus inhibiting recovery of caribou populations (Boulanger et al., 2024; Dickie et al., 2017, 2023; Finnegan et al., 2018; Fullman et al., 2025). Additionally, migratory species need networks of protected habitat that are connected to facilitate their seasonal movements, and they need to encompass areas used in the past and areas expected for use in the future (Gunn et al., 2008; Singh & Milner-Gulland, 2011; Taillon et al., 2012). The DU caribou management plan, released in 2018 prior to the current GBRP proposal, addresses habitat protection; while mining (5.2.7) and roads (5.2.8) were assessed as low impact at the time, the impact of the proposed development plans are likely much greater than thought in consistently used caribou and hunting areas (Campbell et al., 2021; Hanke et al., 2021, 2024; Worthington et al., 2018).

Despite evidence supporting significant impacts to caribou, major development projects are continually licensed under the pretense of sufficient protection or mitigation measures (Cameron & Kennedy, 2023; Collard et al., 2020) or through the strategic use of scale, Inuit knowledge, and consultation efforts (Cameron & Kennedy, 2023). Mitigation measures play a critical role in these assessments but review studies indicate these measures cannot adequately protect caribou from habitat and population loss (Cameron & Kennedy, 2023; Collard et al., 2020) and are frequently resisted or violated post-licensure (Bernauer et al., 2023; Cameron & Kennedy, 2023). It is unlikely that the Endangered DU caribou herd, already at a historic low, can sustain the additional cumulative impacts from the proposed infrastructure nor the mining of critical minerals that will be unlocked afterwards, let alone recover to their former numbers.

### The federal government is lagging behind duties for the Species at Risk Act

The federal decision to uplist the DU caribou as Endangered is nearly six years overdue (COSEWIC, 2017; Government of Canada, 2019b). The Minister of the Environment received the COSEWIC reassessment on 2018-10-15, posted their response on 2019-01-12, and the Governor in Council decision was due by 2020-10-15 (Government of Canada, 2019a). No decision has been issued at the time of this writing. The Nunavut Wildlife Management Board submitted support for the federal uplisting of the herd in 2022, and the Government of Northwest Territories uplisted the herd to Endangered under their territorial Species at Risk Act in July 2024 (Northwest Territories, 2024; Nunavut Wildlife Management Board, 2022). We know that the DU caribou range has contracted as their abundance has dropped, and yet the hotspots we have identified are still used consistently by caribou and are effectively critical habitat for the herd (Campbell et al., 2021; Haniliak et al., 2025; Hanke et al., 2024). Habitat protection afforded under Northwest Territories territorial jurisdiction only covers a fraction of the caribou range (Fig. 1), leaving critical habitat open for development (Species at Risk Committee, 2023; Worthington et al., 2018).

### Inuit harvesting rights will be compromised by the GBRP project

In the context of this study, disrupting these harvesting hotspots will negatively impact food sovereignty by reducing autonomy, will amplify food insecurity through displacement or disappearance of caribou, and will make it harder for Inuit to practice and connect to their culture (Inuit Circumpolar Council Alaska, 2015, 2020). The three hotspots apparent in our results are closely associated with family connections: people often go harvesting where their families had come from before settlement. This connection between caribou and Indigenous social systems extends far beyond these data and this study system (Borish et al., 2021, 2022; Caughey et al., 2024; Gagnon et al., 2023). Understanding the familial background to the data points can help with data interpretation, and these dimensions add meaning to the impact of the results (Gagnon et al., 2023; Hanke et al., 2024). Disrupting harvesting hotspots means losing knowledge of how to safely travel and engage in search and rescue missions, skills increasingly recognized and valued in the context of Canadian Arctic Sovereignty, and is a serious challenge to Inuit harvesting rights.

The GBRP project will have negative effects on caribou conservation and Inuit harvesting rights that are unmitigable in nature: regardless of caribou disappearance or displacement, there will be cumulative impacts on the caribou, and there will be harm to Inuit harvesting rights. These results support the stance taken by the Hunters and Trappers Organizations of Burnside and Kugluktuk in opposition to the GBRP project. The question of “which organization is the legitimate representative of Inuit rights and interests vis-à-vis extractive industries”, and who ultimately grants or withholds Inuit consent to development is of critical importance to the GBRP project (Bernauer, 2025, p. 42; Peres & Stanley, 2025). We suggest the divergence in opinion between the rightsbearing groups should disqualify the GBRP project from streamlined regulatory approval and warrant careful consideration at the territorial level.

### Reviews and licensure should be based on evidence and rooted in cross-legislation obligations

In this paper, we have demonstrated how Indigenous and Western science can come together to empirically identify critical caribou habitat and important Inuit cultural locations. Leading international organizations tasked with biodiversity protection, e.g. the Intergovernmental Science-Policy Platform on Biodiversity and Ecosystem Services or the Convention on Biological Diversity, promote such approaches that cross knowledge systems (Díaz et al., 2015; Reyes-García et al., 2023). In Canada, prioritizing Indigenous knowledge systems to address conservation concerns is a legislative responsibility, e.g. Species at Risk Act, the Nunavut Land Claims Agreement Act, the Impact Assessment Act. Within the development arena, environmental assessments are tasked to come to rational, evidence-based decisions that balance ecological, social, and economic goals; in Canada, they must consider information from Indigenous and Western knowledge systems (Austman & Debicki, 2025; Cameron & Kennedy, 2023; Eckert et al., 2020). Tension often lies between these state responsibilities and advancing economic growth, found inside and outside Canada (M’Gonigle & Takeda, 2013); federally and provincially (Collard et al., 2020); territorially and regionally (Bernauer, 2025; Bernauer et al., 2023; Bernauer & Cameron, 2026; Peres & Stanley, 2025).

Canadian Prime Minister Carney claims that Canada, and other middle power countries, must act consistently, build what they claim to believe in, and remain committed to sustainability (Carney, 2026). Standing by these tenets is incredibly difficult in the complex case of critical minerals and mineral security, comprising conflicting policies related to geopolitical turmoil, the fourth industrial revolution, green energy transition, Indigenous rights, and biodiversity conservation. However, it is important to remember that energy transition based solely on minerals does not have guaranteed feasibility (Henckens, 2022), there is a lack of evidence that stated mitigation measures actually protect the environment (Cameron & Kennedy, 2023), proponent-led review systems favour biased assessments and reporting (Arsenault et al., 2019), and most financial benefits from mining are found to accrue to non-Indigenous equity partners and creditors, mining companies, and their shareholders rather than the local people (Inutiq et al., 2024; Peres & Stanley, 2025). It is crucial that development project reviews and licensure are based on evidence, and not obfuscated data, and are rooted in cross-legislation obligations; this is the essence of Prime Minister Carney’s commitment to act consistently and in line with previously stated values.

## Acknowledgments

We thank the numerous Inuit Knowledge experts with whom we work alongside. Ideas were formed through close collaborations between the broader teams at the Kugluktuk Angoniatit Association, Cloudberry Connections, Kutz Research Group at the University of Calgary, and Lisa-Marie Leclerc with the Government of Nunavut.

## Data availability statement

The datasets presented in this article are not readily available because of the nature of this research and its focus on Indigenous Knowledge. The participants of this study did not consent for their data to be shared publicly beyond what is included in the paper. Requests to access the Inuit Knowledge or sample datasets should be directed to SK, https://vet.ucalgary.ca/groups/arctic-wildlife-health/contact. Requests to access the Government of Nunavut data should be directed to the Department of Environment, Government of Nunavut.

## Positionality statement

The team leading this research has many years of experience in community-based research in Inuit communities. AH has worked with Inuit in Kugluktuk who harvest DU caribou since 2017 to document their knowledge and make it more available and accessible for decision-making. AD is Inuk, a resident of Kugluktuk, and the general manager for the Kugluktuk Angoniatit Association. SK has worked in Kugluktuk since the 1990s on various community-based wildlife health monitoring projects and leads the Kutz Research Group at the University of Calgary. DB has worked with wildlife co-management boards across the Canadian Arctic to co-create visual media about Indigenous Knowledge of species like polar bears, caribou, and beluga. AH completed their PhD from 2017-2024 under the supervision of SK with AD as a thesis advisor, and they now work at Cloudberry Connections with DB.

